# Short RFamide and CFamide peptides as novel positive modulators of Acid-Sensing Ion Channel 3 with similar potentiating effects but different reversibility

**DOI:** 10.1101/2025.07.16.665058

**Authors:** Maurizio Toft, Maëva Meynier, Hélène Lubrano Di Scampamorte, Cédric Vallée, Miguel Salinas, Peijun Zhang, Emmanuel Bourinet, Eric Lingueglia, Emmanuel Deval

**Affiliations:** Université Côte d’Azur, CNRS UMR7275, Inserm U1323, IPMC, LabEx ICST, FHU InovPain, Valbonne 06560, France; Electron Bio-Imaging Centre (eBIC), Diamond Light Source Ltd, Harwell Science and Innovation Campus, Didcot, United Kingdom; Division of Structural Biology, Wellcome Centre for Human Genetics, University of Oxford, Oxford, United Kingdom; IGF, Montpellier University, CNRS, INSERM, Montpellier 34000, France

## Abstract

Acid-sensing ion channels (ASICs) are members of the DEG/ENaC family that includes the only known peptide-gated ion channels. While ASICs are gated by protons, they have kept sensitivity to peptides and are notably modulated by the molluscan FMRFamide and other related mammalian neuropeptides ending by the RFamide motif. Screening efforts have been made to identify and characterize natural peptides able to modulate ASICs’ activity, and some peptides from different species are already known to modulate ASIC1a, ASIC1b and ASIC3. We identified here a set of synthetic short amidated hexapeptides, initially designed thirty years ago for their ability to inhibit the Na/Ca exchanger, as potent and selective positive modulators of the ASIC3 acid-induced activity. We focused on two of them, a RFamide peptide (FR<u>CC</u>RFamide) and a CFamide peptide (FRC̄R̄C̄Famide), demonstrating that they have similar specificity for and effects on ASIC3. The potentiating effects of the two peptides are due to a strong slow-down of the current desensitization, leading to an increase in the amount of current induced by acid pH (≤pH6.6), with apparent affinities ranging from 1 to 5 µM. Surprisingly, the washout kinetic for the FRC̄C̄RFamide peptide was much slower than those of FRC̄R̄C̄Famide and other known RFamide peptides, suggesting potential differences in their mechanisms of action. Computational modeling and structure-function analysis reveal interactions of both peptides with the non-proton binding site of ASIC3 initially identified for the synthetic compound GMQ (2-guanidine-4-methylquinazoline), as already reported before for other RFamide peptides, but our data also suggest possible additional effects of FRC̄C̄RFamide involving directly or indirectly the proton binding domain. These findings expand our understanding of peptide modulation of ASIC channels and identify novel pharmacological tools selective among ASICs for investigating ASIC3 function.

## Introduction

Several members of the degenerin/epithelial sodium channel family of ion channels (DEG/ENaC) have the interesting property of being directly modulated by peptides (for review, see (1)). Some invertebrate members of this ion channel family (*i.e.*, FaNaCs (2) from snails, HyNaCs from hydra (3) and MGIC from a marine annelid (4)) even represent the only known peptide-gated ion channels, as they are directly opened by RFamide (FaNaCs and HyNaCs) or Wamide (MGIC) peptides. Mammalian Acid-Sensing Ion Channels (ASICs) also belong to the DEG/ENaC family. They are gated by extracellular proton and display both transient and sustained depolarizing activities triggered by extracellular acidification (for reviews see (5, 6)), and also alkalization in the case of human ASIC3 (7). Although they are not peptide-gated ion channels, ASICs have kept sensitivity to peptides, as they are positively modulated by the canonical molluscan FMRFamide and other endogenous RFamide-related neuropeptides, which mainly target ASIC3, by dynorphin A and big dynorphin targeting ASIC1a, and by YFMRFamide that modulates ASIC1b and ASIC3 (8–14). While the existence of endogenous peptide(s) able to directly gate ASICs remains an open question, no endogenous peptide activator has been found so far despite extensive search.

RFamide peptides are major effectors of ASICs, mainly ASIC3, which is positively modulated by FMRFamide, neuropeptide FF (NPFF), neuropeptide SF (NPSF), YFMRFamide and RPRFamide, with EC_50_ ranging from 2 to 50µM (8–14). The RFamide motif at the end of these peptides appears very important for their effects, *i.e.* adding this motif to Met-enkephalin that has no effect on ASIC3 is sufficient to produce an effect comparable to that of FMRFamide (11), although RFamide alone remains much less efficient than FMRFamide, NPFF or NPSF (10). More recently, a RPRFamide peptide isolated from *Conus textile* venom has been reported to target ASIC3, enhancing its activity as well as acid-induced muscle pain (13). This cono-RFamide peptide has been proposed to stabilize the open conformation of ASIC3 through binding to its non-proton binding domain (12), which could be a common mechanism for all RFamide peptides.

About thirty years ago, seven positively-charged cyclic amidated hexapeptides were described as novel blockers of the cardiac sodium-calcium exchange (NCX), with IC_50_ ranging from 10µM to more than 250µM (15). Part of these synthetic peptides display a C-terminal RFamide motif, while others display a C-terminal CFamide motif, raising the question of whether these peptides, including the non-RFamide ones, can also modulate ASICs, particularly ASIC3. Here, we report that these peptides are indeed novel positive modulators of ASIC3 and we focus on two of them: *(i)* the FRC̄R̄C̄Famide, which is not a RFamide peptide and has the best NCX inhibitory activity (15) and *(ii)* the FRC̄C̄RFamide, which is a RFamide and has a 15-fold lower inhibitory effect on NCX (15). Both peptides display a similar potentiating effect on ASIC3, with EC_50_ in the micromolar range and interaction sites within the non-proton binding site of the channel initially identified for the synthetic compound GMQ (2-guanidine-4-methylquinazoline) (16). This is in line with what has already been proposed for other RFamide peptides (12). However, the washout kinetic for the FRC̄C̄RFamide peptide was much slower than those of FRC̄R̄C̄Famide and other RFamide peptides, suggesting a more complex mechanism of action also possibly involving the proton binding domain.

## Materials and methods

### Plasmid constructions and mutagenesis

The coding sequences of rat ASIC1a, ASIC3 and human ASIC3a and their related mutants were subcloned into pIRES_2_-eGFP vector (Addgene). Rat ASIC1b, ASIC2a and ASIC2b were subcloned into pCI vector (Promega). Mutants of rat ASIC3 were obtained by recombinant PCR strategies as previously described (17).

### Cell culture and transfection

HEK293 cell were grown in DMEM medium (Lonza™ BioWhittaker™) supplemented with 10% of heat-inactivated fetal bovine serum (<u>Biowest</u>^TM^) and 1% of antibiotics (penicillin + streptomycin, Lonza™ BioWhittaker™). One day after plating, cells were transfected with either pIRES2-ratASIC1a-EGFP, pCI-ratASIC1b + pIRES2-eGFP, pCI-ratASIC2a + pIRES2-eGFP, pCI-ratASIC2b + pIRES2-eGFP, pIRES2-ratASIC3-EGFP, pIRES2-humanASIC3a-EGFP vector using the JetPEI reagent according to the supplier’s protocol (Polyplus transfection SA, Illkirch, France). Fluorescent cells were used for patch-clamp recordings 2–4 days following transfection.

### Primary culture of DRG neurons

Lumbar dorsal root ganglia (DRG L1 to L6) were rapidly dissected out from euthanized adult male C57Bl6J (Janvier Lab, at least 8 weeks of age) and ASIC3^-/-^ mice, and placed in cold Ca^2+^-free/Mg^2+^-free HBSS medium (Corning^®^) supplemented with 10mM glucose, and 5mM HEPES (pH7.5 with NaOH). After carefully removing the roots, DRGs were enzymatically dissociated at 37°C for 2 x 20min in the HBSS solution supplemented with calcium (CaCl_2_ 5 mM) and containing type II collagenase (Gibco, France) and dispase (Gibco, France). DRGs were then mechanically triturated and washed in a complete neurobasal A medium (NBA, Gibco^®^ Invitrogen^TM^ supplemented with 2% B27, 2mM L-glutamine and 1% antibiotics: penicillin + streptomycin, Lonza™ BioWhittaker™), before being plated in 35-mm petri dishes. Neurons were then kept in culture at 37°C for one week with 1/3 of the culture medium (complete NBA supplemented with 10ng/ml NT3, 2ng/ml GDNF, 10ng/ml BDNF, 100ng/ml NGF and 100nM retinoic acid) renewed every 2 days, and were used for patch-clamp experiments at least 1 day after plating.

### Patch-clamp experiments

Membrane currents were recorded using patch-clamp in the whole cell configuration (voltage-clamp mode). Patch pipettes (3-7MΩ) were made from borosilicate glass tubes (Hilgenberg, Germany) using a P30 puller (Sutter instruments, USA), before being filled with an intracellular solution containing either 135 mM KCl, 2.5mM ATP-Na_2_, 2.1 mM CaCl_2_, 5 mM EGTA, 2 mM MgCl_2_, 10 mM HEPES (pH7.3 with KOH) for DRG neurons, or 135 mM KCl, 5mM NaCl, 5 mM EGTA, 2 mM MgCl_2_, 10 mM HEPES, (pH7.3 with KOH) for HEK293 cells. Cells were bathed into an extracellular solution made of 145mM NaCl, 5mM KCl, 2mM CaCl_2_, 2mM MgCl_2_ and 10mM HEPES (or MES depending on the final pH of the solution), and this medium was supplemented with 10mM glucose for DRG neurons. Extracellular pH was then adjusted to its final value using NMDG, including HEPES-containing solutions for 7.4≤pH≤6.6 and MES-containing solutions for 6.6<pH≤5.0. During patch-clamp recordings, the cell under investigation was continuously superfused with extracellular solutions using a homemade eight-outlet system digitally controlled by solenoid valves (Warner Instruments), allowing rapid changes of the immediate cell environment from control to test solutions. Electrophysiological signals generated by patch-clamped cells were amplified and low-pass filtered at 2kHz using an Axopatch 200B amplifier (Molecular Devices, UK), digitized with a 1440 A-D/D-A converter (Molecular Devices, UK), sampled at 20kHz and stored on a computer using Clampex software (V10.7, Molecular Devices). Off-line analyses were then made using Clampfit software (V10.7, Molecular Devices).

### Chemicals

All the peptides used in this study were synthetized by the SB Peptides company (SB Peptides, France). They were all prepared as 1 mM stock solutions in water or saline (extracellular patch clamp medium) and stored at -20°C until their use at final concentrations for patch-clamp experiments (1nM to 100 µM). In order to avoid peptides adsorption to the perfusion tubing, BSA 0.05% was added to the resting pH7.4 extracellular patch-clamp solution.

### Computational methods

The 3D structures of the homotrimeric ASIC3 was predicted using AlphaFold3 (18) (https://alphafoldserver.com/) using the *Rattus norvegicus* ASIC3 sequence from Uniprot (entry O35240). Residues 495 to 533 (C-ter) were removed from the structure due to low pLDDT scores (less than 50) and lack of secondary structure. The resulting rASIC3 structure was then embedded in a POPC (1-palmitoyl-2-oleoyl-sn-*glycero*-3-phosphocholine) bilayer membrane, solvated in water with the TIP3P model (19), and neutralized with 0.15 M of NaCl using the CHARMM-GUI Membrane Builder input generator (20). The system was subsequently minimized using the steepest decent algorithm with position restraints on the heavy atoms of the protein and the lipids, then was equilibrated by 6 equilibration steps following the recommendation from CHARMM-GUI. A short Molecular Dynamics simulation was performed for 10 ns on the equilibrated system and a cluster analysis was performed using the ‘gmx cluster’ command from GROMACS to extract the most representative conformation of the embedded rASIC3 (referred to as the ‘post-MD’ structure). The 3D structures of both peptides (FRC̄R̄C̄Famide and FRC̄C̄RFamide) were generated using the ‘Build Structure’ tool from ChimeraX version 1 (21). Docking of both peptides was then performed on the extracellular domain (search box size of 100 Å x 100 Å x 100 Å) of both the AlphaFold structure and the post-MD structure from the cluster analysis using the AutoDock Vina extension (version 1.1.2) of Chimera version 1.17.3 (22, 23). Further docking of both peptides (search box size of 25 Å x 25 Å x 25 Å) were performed on the two sites with the most occupancies (determined from the results of the extracellular docking) referred to as the ‘proton site’ and the ‘non-proton site’. As rASIC3 is an homotrimer, 3 dockings per site and per peptide were performed. The best docking results (*i.e*., with the lowest scores) for both peptides were used to generate new systems containing the peptide in one of the two sites of rASIC3 embedded in a POPC membrane using CHARMM-GUI, as previously described. The CT2 terminal group patching was used for the peptides to make sure they have the amide functional group at their C-ter. Then, 3 independent MD replicates of each system with either peptide in the different docking sites were produced for 30 ns, for a total of 2,160 ns of simulation. Finally, after discarding the first 5 ns of the simulations, the trajectories of each run were further analyzed using ProLIF (Protein-Ligand Interaction Fingerprints) (24) to identify interactions between the peptide and the channel and the ‘gmx energy’ command to quantify the interaction energies for all residues interacting with the peptides for more than 50% of the simulation in at least one of the replicates. MD production runs were performed in the NPT ensemble using the V-rescale thermostat and the C-rescale barostat to maintain the temperature and the pressure at 310.15 K and 1 bar, respectively. The leap-frog integrator was used with a timestep of 2 fs, and all covalent hydrogen bonds were constraint with the LINCS algorithm (25). The cutoff for the short range non-bonded interactions was 1.2 nm, and the long-range electrostatic interactions were evaluated using the Particle Mesh Ewald (PME) methods (26). All simulations were carried with GROMACS version 2024.1 with CUDA GPU support, and the CHARMM36m forcefield (27) (28).

### Statistical analysis

Data are presented as mean ± SEM and statistical analyses were performed using GraphPad Prism software. Differences between sets of data were considered significant when *p*-values were less or equal to 0.05. Significant differences between datasets calculated from either parametric or non-parametric tests followed by adequate *post-hoc* tests (see figure legends).

## Results

### The cyclic FRC̄R̄C̄Famide peptide is a selective potentiator of ASIC3 current

Among the cyclic hexapeptides initially synthetized as inhibitors of NCX activity, FRC̄R̄C̄Famide was reported as the one with the best affinity (IC_50_ ∼10µM, see (15)). Although this peptide does not contain the final RFamide motif, we found it to be a strong and quite selective potentiator of ASIC3 activity evoked at pH5.5 in HEK293 transfected cells (**Fig.1**). This effect was characterized by a typical slow-down of the current desensitization when the peptide was applied at 10µM in the resting/conditioning pH7.4, leading to an increase of its pH5.5-evoked sustained component (see the sustained current amplitude measured 5 sec after the current onset, **Fig. 1A**), with no significant effect on the transient peak current (**Fig. 1A-B**). Conversely, this effect was not observed when the peptide was applied in the acid test pH5.5 solution (**Fig. 1A-B**). Finally, co-applying the peptide both in the pH7.4 and pH5.5 extracellular solutions did not induce any supplemental effect compared to application at pH7.4 only (**Fig. 1A-B**). The potentiating effect of FRC̄R̄C̄Famide on pH5.5-evoked ASIC3 sustained activity was highly variable (between 1.9- and 41.8-fold increase, **Supplementary** Fig. 1A), and it was negatively correlated to the basal pH5.5-sustained current amplitude evoked in control condition, *i.e.* without peptide. Indeed, the strongest effects of FRC̄R̄C̄Famide (≥20-fold increase) were observed when the absolute value of the basal pH5.5-evoked sustained current amplitudes were the smallest (-0.5 to -6.0 pA/pF), and FRC̄R̄C̄Famide effect potency decreased as the absolute value of the basal pH5.5-evoked sustained current amplitudes increased (**Supplementary** Fig. 1A).

**Figure 1:**
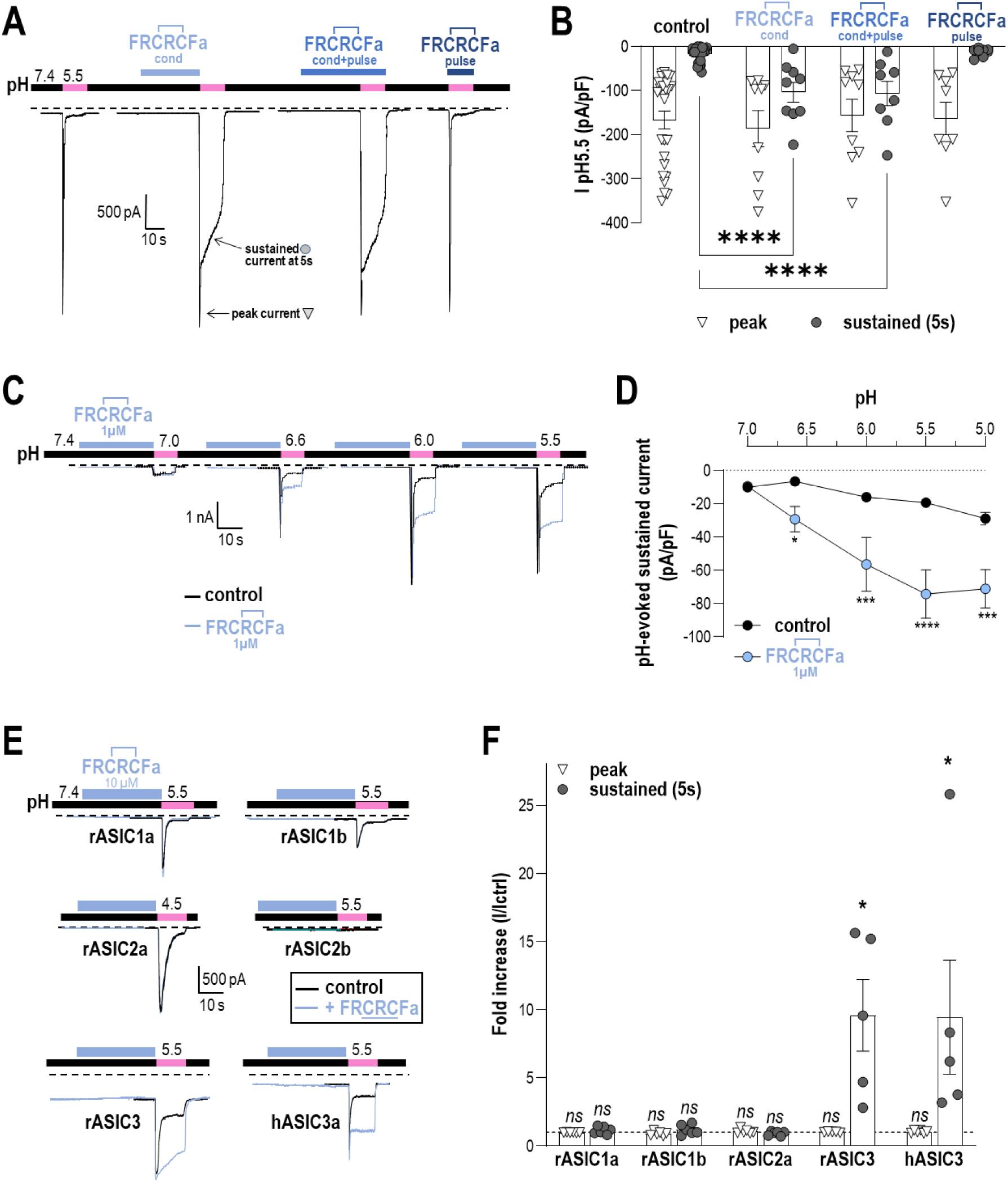
FRC̄R̄C̄Famide acts on the close state of ASIC3 to potentiate its low acid pH-induced sustained activity. ***A-B*,** Typical effects of FRC̄R̄C̄Fa peptide applied extracellularly at 10 µM on pH5.5-evoked rat ASIC3 currents recorded at -50mV in HEK293 transfected cells (**A**). FRC̄R̄C̄Fa peptide (10µM) was either applied into the pH7.4 resting solution (conditioning, cond), the test pH5.5 solution (pulse) or both (cond + pulse, n=8-25; ****p<0.0001, mixed effect analysis followed by a Dunnett multiple comparison test) (**B**). ***C-D,*** Effect of FRC̄R̄C̄Fa applied at 1µM in the pH7.4 resting solution on ASIC3 current evoked by different pH (n=4, *p>0.05, ***p<0.001 and ****p<0.0001, Two-way ANOVA followed by a Sidak’s multiple comparison test). ***E*,** Typical effects of FRC̄R̄C̄Fa (10µM) on homomeric rASIC1a, rASIC1b, rASIC2a, rASIC2b, rASIC3 and hASIC3a currents. ***F,*** Fold increase factors of the peak and sustained currents (Ipeptide/Icontrol, I/Ictrl) generated by FRC̄R̄C̄Fa (10µM) on the different ASICs shown in E (n=4-6, *p<0.05, paired *t*-test). As no significant pH5.5-evoked current was recorded for ASIC2b with or without peptide, the fold increase factor is not represented for this channel (n=4).

The effect of FRC̄R̄C̄Famide was also observed when the peptide concentration was reduced to 1µM (**Fig. 1C-D**), with still a significant potentiation of the pH-evoked ASIC3 sustained activity while its transient peak amplitude remained unaffected (**Supplementary** Fig. 1B). The potentiation was strongest when ASIC3 sustained currents were generated by relatively more acidic pH (≤ pH6.6), *i.e.*, outside the ASIC3 window current (29–31).

The FRC̄R̄C̄Famide peptide was next tested on different ASIC homomeric channels expressed in HEK293 cells (**Fig. 1E-F**), including ASIC1a, ASIC1b, ASIC2a and ASIC2b, in addition to ASIC3. Except for ASIC3 (both rat and human clones), none of the other ASIC homomeric channels had their acid-evoked currents potentiated when FRC̄R̄C̄Famide (10µM) was applied extracellularly in the resting/conditioning pH7.4, showing a relatively good selectivity toward ASIC3 compared to other ASICs, at least at the tested concentration of 10µM (**Fig. 1E-F**).

### The cyclic FC̄R̄C̄RFamide and FRC̄C̄RFamide peptides also potentiate ASIC3 current

We next compared the effects on ASIC3 of the FRC̄R̄C̄Famide peptide with those of another cyclic hexapeptide ending by the RFamide motif, *i.e.* FRC̄C̄RFamide (**Fig. 2**), which has a relatively low IC_50_ for NCX (150µM, see (15)). The two peptides were found to have a similar effect, *i.e.*, a drastic potentiation of the pH5.5-evoked sustained ASIC3 current when applied at 10µM in the resting pH7.4, with no significant effect on the peak current amplitude (**Fig. 2A-B**). Because these effects were associated to a slowdown of desensitization, a precise analysis of this current phase was performed with and without peptides (**Fig. 2C**). There was no significant difference between the effects of the two peptides, and similar effects were also found with another cyclic hexapeptide ending by the RFamide motif, *i.e.* FC̄R̄C̄RFamide (**Supplementary** Fig. 2). Overall, both the CFamide and RFamide cyclic hexapeptides displayed similar potentiating effects on ASIC3, and we next focused our investigations on two of them: FRC̄R̄C̄Famide and FRC̄C̄RFamide.

**Figure 2:**
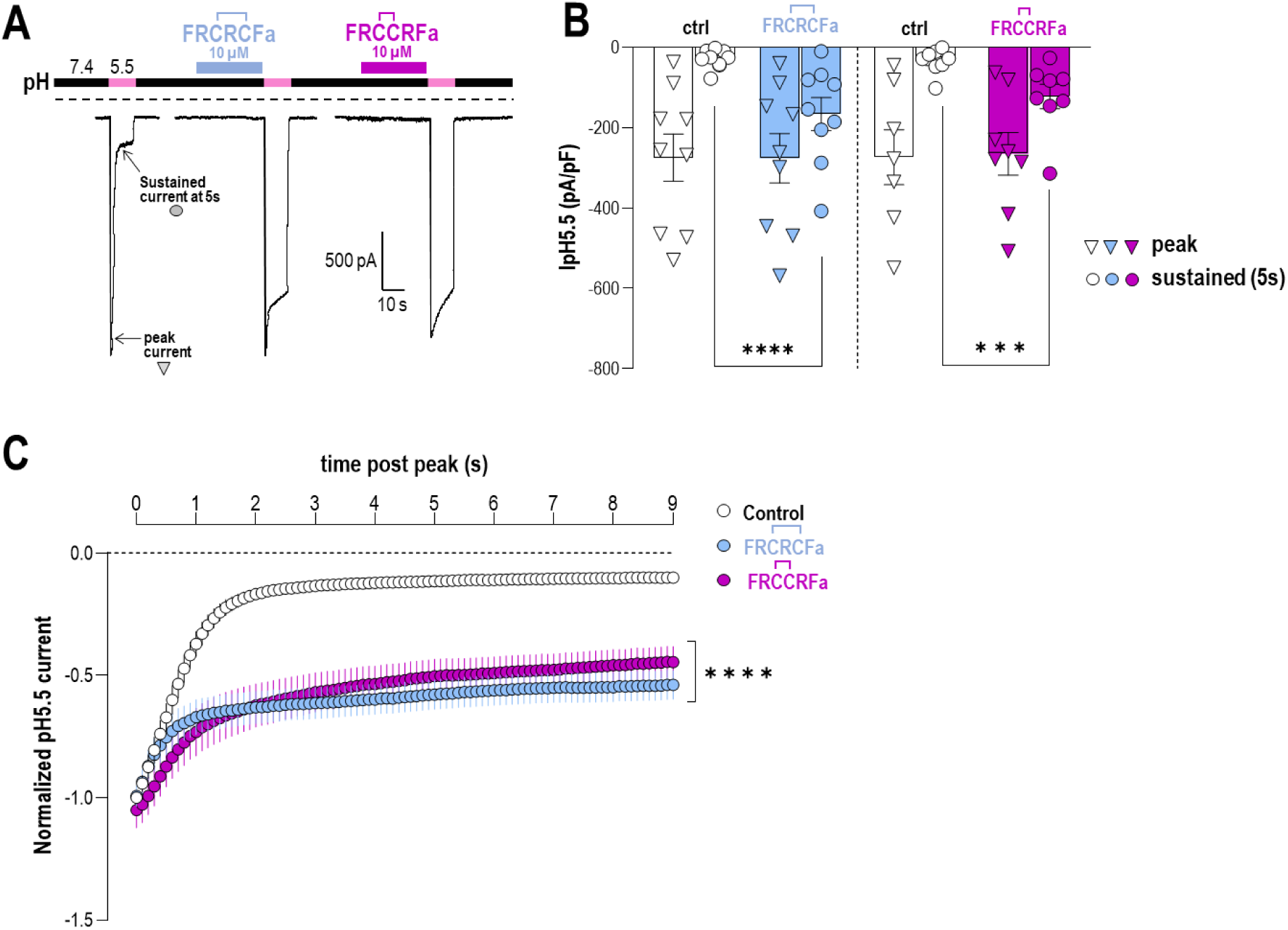
Cyclic CFamide and RFamide peptides similarly potentiate ASIC3 sustained current. ***A,*** Typical effect of FRC̄R̄C̄Famide and FRC̄C̄RFamide peptides (10µM in the pH7.4 resting solution) on rat ASIC3 activity. Currents were generated by pH drops from resting pH7.4 to pH5.5 and were recorded at –50 mV in HEK293 transfected cells. ***B,*** Analysis of the peak and sustained current densities elicited at pH5.5 before and after extracellular applications of peptides (sustained currents were measured 5s after initiation of ASIC3 currents, n=9 and 7 for FRC̄R̄C̄Famide and FRC̄C̄RFamide respectively; ***p<0.001 and ****p<0.0001, Mixed-effect analysis followed by Dunnett’s multiple comparisons test). ***C,*** Analysis of ASIC3 current desensitization at pH5.5. Current amplitudes were measured every 100ms during the 9s following the peak, and normalized to the control peak amplitude before application of each peptide (n=13, 9 and 8 for control, FRC̄R̄C̄Famide and FRC̄C̄RFamide respectively; ****p<0.0001 compared to control, Mixed-effect analysis followed by Tukey’s multiple comparisons test).

### FRC̄R̄C̄Famide and FRC̄C̄RFamide have different washout kinetics

The on/off kinetics of the effects of both FRC̄R̄C̄Famide and FRC̄C̄RFamide on ASIC3 (**Fig. 3**) were analyzed by first varying the application time of each peptide (10µM in pH7.4 resting solution, **Fig. 3A**). We found that the onset of FRC̄C̄RFamide effect was very similar to that of FRC̄R̄C̄Famide, with a half maximum effect reached in 1-2 seconds and a maximal effect in ∼20 seconds (**Fig. 3B**, τ_onset_ =3.8s for both FRC̄R̄C̄Famide and FRC̄C̄RFamide). Analysis of the washout of FRC̄R̄C̄Famide and FRC̄C̄RFamide effects (**Fig. 3C**) highlighted a slower off kinetic (τ_off_) for FRC̄C̄RFamide (4-5 times slower than FRC̄R̄C̄Famide, **Fig. 3D**, τ_off_ =11.9s and 65.5s for FRC̄R̄C̄Famide and FRC̄C̄RFamide, respectively), pointing out the poor reversibility of this peptide’s effect. Interestingly, the slow washout kinetic of FRC̄C̄RFamide was drastically accelerated by increasing the frequency of pH5.5 pulses during the washout protocol, whereas that of FRC̄R̄C̄Famide remained unchanged (τ_off_ =22.7s and 13.1 for FRC̄C̄RFamide and FRC̄R̄C̄Famide, respectively, see **Supplementary** Fig. 3), indicating more complex interactions between ASIC3 and the FRC̄C̄RFamide peptide. Finally, we performed dose-response studies for both FRC̄R̄C̄Famide and FRC̄C̄RFamide peptides (**Fig. 3E**), and found EC_50_ values of 2.87±0.44µM and 0.98±0.16µM, respectively, indicating a similar range of apparent affinities for ASIC3.

**Figure 3:**
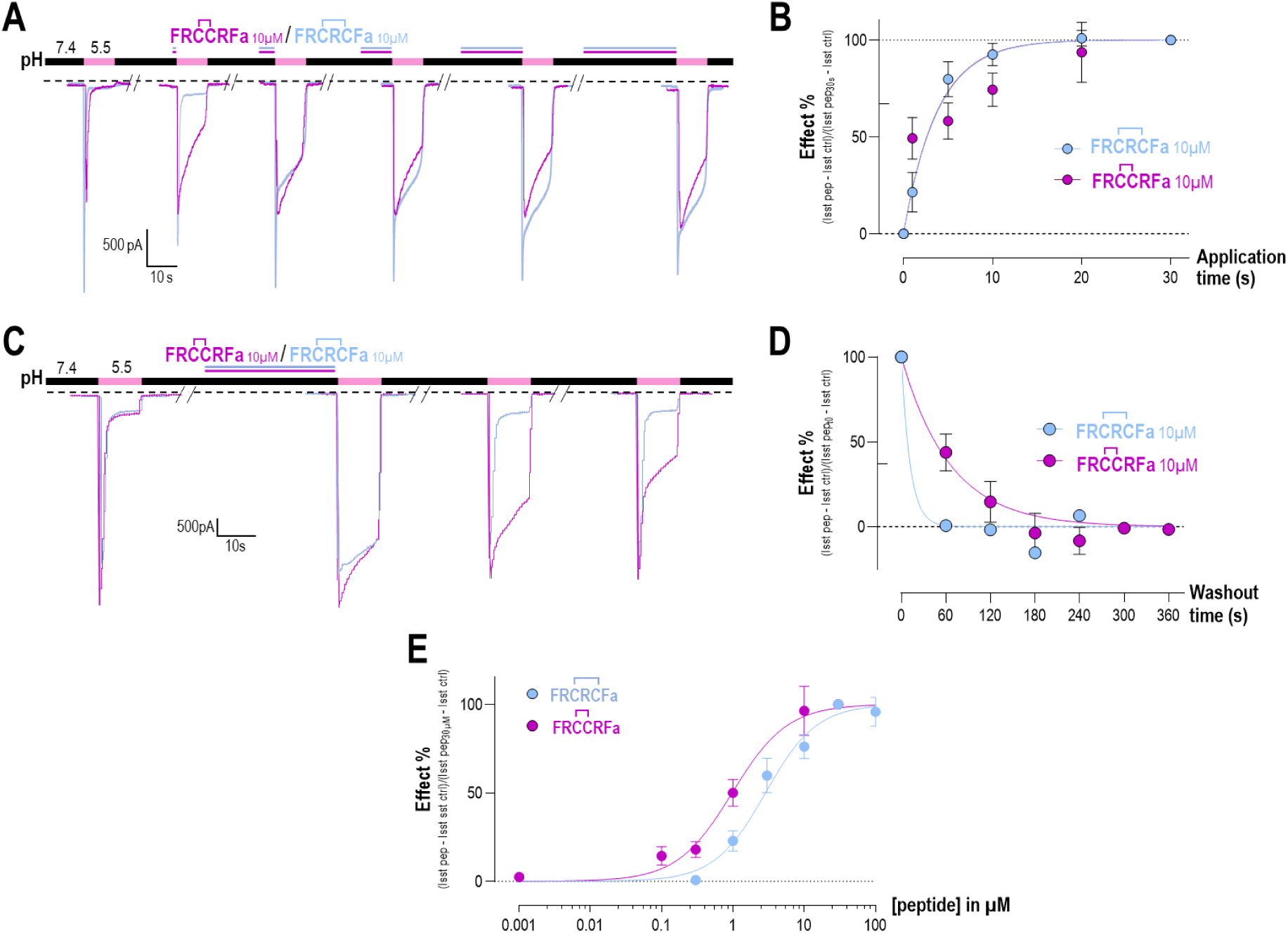
Comparison of the effects and FRC̄R̄C̄Famide and FRC̄C̄RFamide on ASIC3. ***A-B*,** Effect of FRC̄R̄C̄Famide or FRC̄C̄RFamide applied extracellularly for different duration (1 to 30s) at 10µM in the pH7.4 resting solution. Onset kinetics of both peptides (B) were fitted over 30s using a one-phase exponential (τ_on_=3.8s for both FRC̄R̄C̄Famide or FRC̄C̄RFamide, respectively; n=9-13). ***C-D,*** Washout time analysis of the effects of FRC̄R̄C̄Famide and FRC̄C̄RFamide applied at 10µM for 30s in the pH7.4 resting solution. The pH5.5 drops during the washout protocol were applied every 1 min. Off kinetics of both peptides (D) were fitted over 360s using a one-phase exponential (τ_off_=11.9s and 65.5s for FRC̄R̄C̄Fa or FRC̄C̄RFamide, respectively; n=13-18). ***E*,** Dose-response curves for the effects of FRC̄R̄C̄Famide and FRC̄R̄C̄Famide using a sigmoidal curve (n=8-21).

### Linearizing FRC̄R̄C̄Famide and FRC̄C̄RFamide only affects the washout kinetic of FRC̄R̄C̄Famide

The linearized counterparts of FRC̄R̄C̄Famide and FRC̄C̄RFamide, *i.e.* FRCRCFamide and FRCCRFamide without disulfide bonds, were found to have similar abilities to potentiate the pH5.5-evoked ASIC3 sustained activity (**Fig. 4A**), without affecting the transient peak current amplitude (**Fig. 4A, inset**). Moreover, the dose response relationships obtained with these linearized peptides also indicated EC_50_ values in the micromolar range (4.41±0.95µM and 4.73±0.80µM for FRCRCFamide and FRCCRFamide, respectively, **Fig. 4B**), as for their cyclic counterparts (2.17±0.91µM and 1.12±0.20µM, respectively, see **Fig. 3E**). Although linearization appeared to slightly decrease the apparent affinities of the peptides for ASIC3 (EC_50_ shifted form 1.12±0.20µM to 4.73±0.80µM for FRC̄C̄RFamide and FRCCRFamide, respectively, and from 2.17±0.91µM to 4.41±0.95µM for FRC̄R̄C̄Famide and FRCRCFamide, respectively) all peptide EC_50s_ remained within a similar concentration range (1-5µM). The main difference between cyclic and linearized peptides was found on the washout kinetics of FRC̄R̄C̄Famide and FRCRCFamide (**Fig. 4C**). Indeed, although the slow washout kinetic of FRC̄C̄RFamide was conserved in the linearized form of this peptide (τ_off_=61.5 and 69.3s, respectively), the washout kinetic of FRC̄R̄C̄Famide became much slower for FRCRCFamide (τ_off_=12.4 and 84.6s, respectively) (**Fig. 4C**). The slow washout kinetic appears to be characteristic of FRC̄C̄RFamide, FRCCRFamide and FRCRCFamide, but not of FRC̄R̄C̄Famide, nor other previously described tetrapeptides such as FMRFamide and RPRFamide (τ_off_=12.4 and 13.9s, respectively) (**Fig. 4C, inset**).

**Figure 4:**
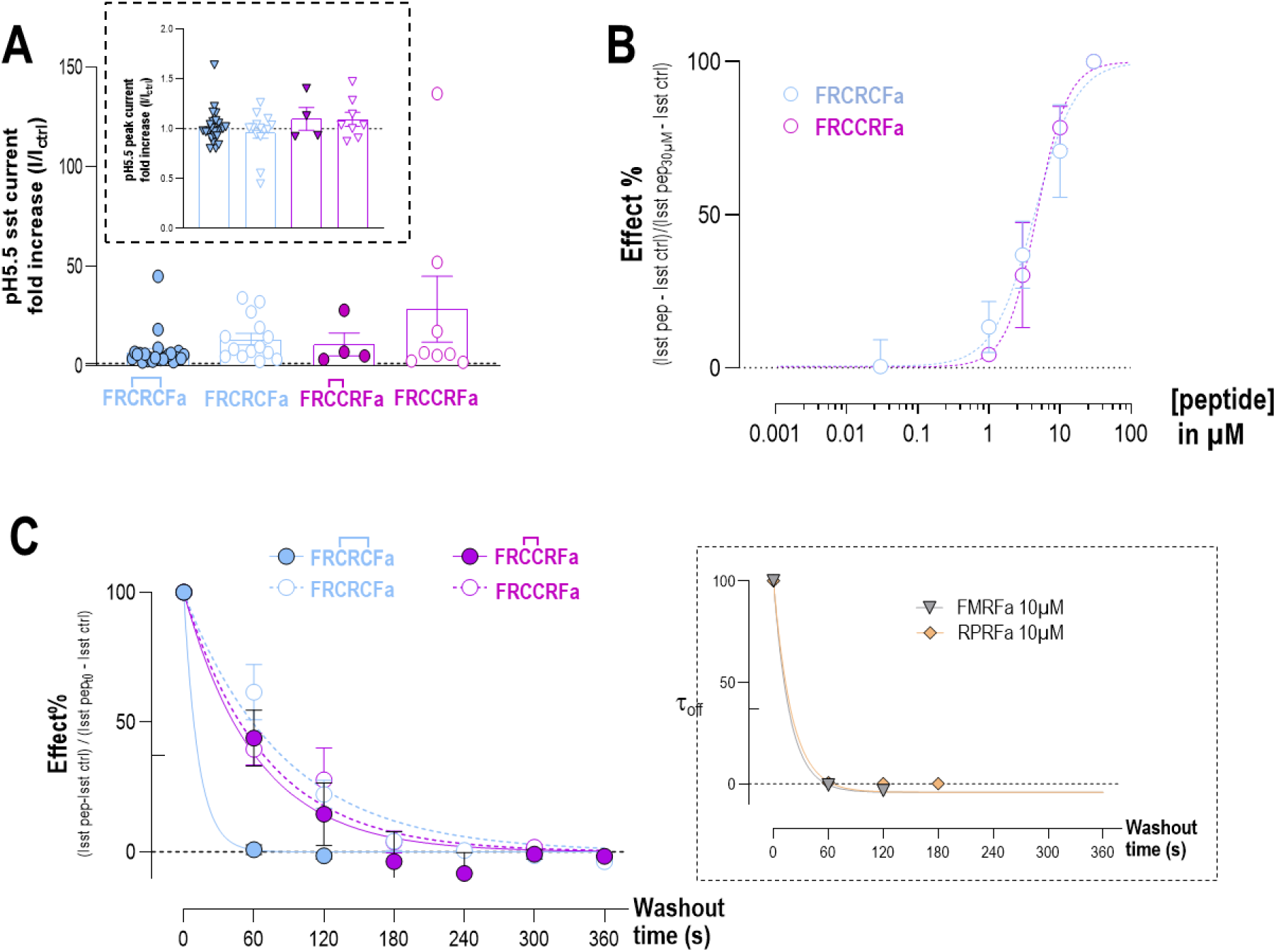
Comparison of the effects on ASIC3 of FRC̄R̄C̄Famide and FRC̄C̄RFamide and of their linearized analogs. ***A,*** Abilities of FRC̄R̄C̄Famide, FRCRCFamide, FRC̄C̄RFamide and FRCCRFamide (10µM) to potentiate the pH5.5-evoked sustained current without affecting the transient peak current amplitude (***inset***) (n=4-24 cells). ***B,*** Dose-response curves of linearized FRCRCFamide and FRCCRFamide peptides using a sigmoidal curve. ***C,*** Washout kinetics of cyclic (FRC̄R̄C̄Famide τ_off_=12.4s and FRC̄C̄RFamide τ_off_=61.5s) and linearized peptides (FRCRCFamide τ_off_=84.6s and FRCCRFamide τ_off_=69.3s) (10µM), and of tetrapeptides FMRFamide (τ_off_=12.4s) and RPRFamide (τ_off_=13.9s) (10µM, ***inset***) (n=3-18 cells).

### FRC̄R̄C̄Famide and FRC̄C̄RFamide specifically potentiate the native pH5.5-induced ASIC3 current in DRG sensory neurons

The effects of FRC̄R̄C̄Famide and FRC̄C̄RFamide peptides were next tested on native acid-evoked currents recorded from primary culture of mouse DRG neurons (**Fig. 5**). Both wild-type (WT) and ASIC3^-/-^ (A3KO) mice were used to perform DRG neuron primary cultures, and peptides were applied onto neurons at 10µM in the pH7.4 resting solution, for 30 seconds before extracellular acidification drops to pH5.5 (**Fig. 5A and D**). Overall analysis of the FRC̄R̄C̄Famide and FRC̄C̄RFamide effects on pH5.5-evoked native currents (I_peptide_/I_ctrl_) clearly showed two populations of neurons in WT DRG (**Fig. 5B**, dotted areas), but not in A3KO DRG (**Fig. 5E**). Indeed, 7 out of 15, and 5 out of 11 neurons from WT DRG had their pH5.5-evoked sustained current strongly potentiated by FRC̄R̄C̄Famide and FRC̄C̄RFamide, respectively (**Fig. 5B-C**), with also some smaller effects on the pH5.5-evoked peak current (**Fig. 5B-C**). The potentiated currents clearly displayed ASIC-like kinetics, with both transient peaks followed by sustained activities of varying amplitudes (**Fig. 5A**, upper panel). Importantly, the potentiating effects of FRC̄R̄C̄Famide and FRC̄C̄RFamide on these ASIC-like currents were lost in ASIC3^-/-^ DRG neurons (**Fig. 5D-F**), demonstrating the involvement of ASIC3. These results further confirm the selectivity of FRC̄R̄C̄Famide and FRC̄C̄RFamide toward ASIC3 among the different ASIC channel subtypes, as shown before on recombinant channels expressed in HEK293 cells (**Fig. 1E**), with potentiating effects observed on the pH5.5-evoked ASIC3 sustained activity. Moreover, the fact that both peptides are active on native ASIC3-type channels also suggests effects on heteromeric ASIC3 channels, which are likely to be functionally expressed in rodent DRG neurons (29, 32). On the other hand, FRC̄R̄C̄Famide and FRC̄C̄RFamide had no effect on WT DRG neurons showing TRPV1-like pH5.5-evoked sustained currents (**Fig. 5A**, lower panel).

**Figure 5:**
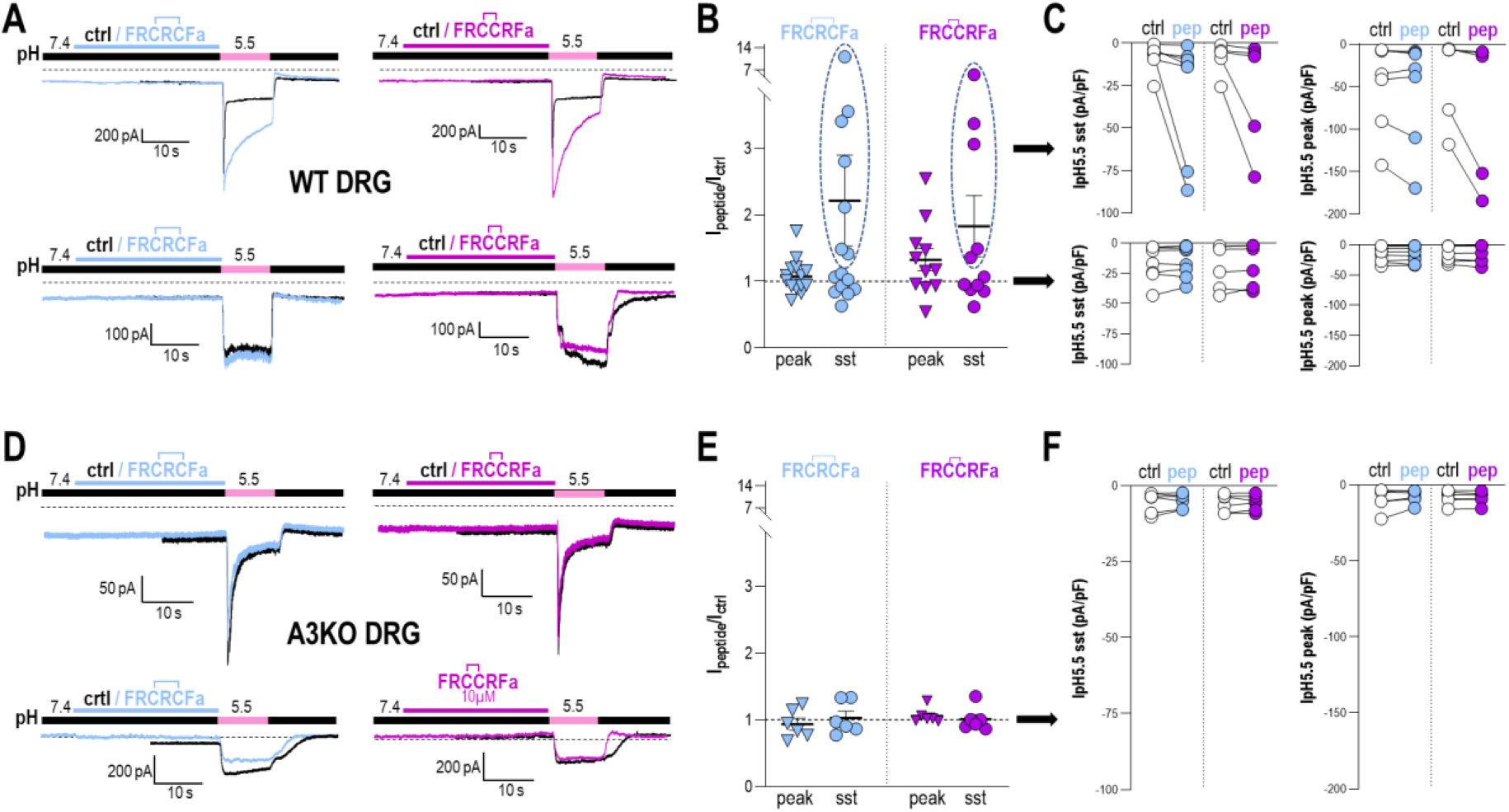
Effects of FRC̄R̄C̄Famide and FRC̄C̄RFamide on native pH5.5-evoked currents recorded in mouse DRG neurons. ***A,*** Typical effects of FRC̄R̄C̄Famide and FRC̄C̄RFamide peptides applied individually at 10µM in the resting pH7.4 extracellular solution on different wild-type (WT) mouse DRG neurons in primary culture. Native pH5.5-evoked currents were recorded at -60 mV, according to the protocol shown by color bars above each current trace. Dotted lines indicate the zero-current level. ***B,*** Fold increase factors (I_peptide_/I_ctrl_) induced by FRC̄R̄C̄Famide and FRC̄C̄RFamide on the peak and sustained currents evoked at pH5.5 in WT DRG neurons. Two populations of neurons can be distinguished, including populations in which peptides have potentiating effects (dotted areas), and another population where they have no effect (data from 15 DRG neurons). ***C,*** Peak and sustained current amplitudes (pA/pF) induced in WT neurons at pH5.5, before and after applications of FRC̄R̄C̄Famide or FRC̄C̄RFamide. Neurons are classified depending on whether the peptides had an affect (upper panel) or not (lower panel), as indicated by the dotted areas in C (see black arrows). ***D,*** Typical effects of FRC̄R̄C̄Famide and FRC̄C̄RFamide applied similarly as in A on ASIC3^-/-^ (A3KO) mouse DRG neurons. ***E,*** Fold increase factors (I_peptide_/I_ctrl_) induced by FRC̄R̄C̄Famide and FRC̄C̄RFamide on the native pH5.5-evoked currents recorded from cultured A3KO neurons (data from 6 neurons). Contrary to WT neurons (B), only one population of neurons was found with no effect of peptides. ***F,*** Peak and sustained current amplitudes (pA/pF) induced in A3KO neurons at pH5.5, before and after applications of FRC̄R̄C̄Famide of FRC̄C̄RFamide.

### In silico structural analysis and site-directed mutagenesis predict interactions of FRC̄R̄C̄Famide and FRC̄C̄RFamide with both the proton and non-proton binding sites of ASIC3

As FRC̄R̄C̄Famide and FRC̄C̄RFamide are both strong potentiators of the ASIC3 sustained activity, the effect of their combined application has then been tested (**Fig. 6A**). When co-applied at concentrations of 5µM, FRC̄R̄C̄Famide and FRC̄C̄RFamide generated a pH5.5-evoked sustained current modulation similar to that observed when the peptides were applied alone at 10µM (**Fig. 6A**), suggesting interactions with a similar binding site. In line with these data, analysis of the washout kinetics revealed that the combination of FRC̄R̄C̄Famide and FRC̄C̄RFamide led to an intermediate effect between those of each peptide applied alone (10µM) (**Fig. 6B**), in good agreement with the hypothesis of a common binding site for both peptides. *In silico* structural modeling was next performed to further study and predict putative interaction sites between the rat ASIC3 channel and the FRC̄R̄C̄Famide and FRC̄C̄RFamide peptides (**Fig. 6C**). The docking predicted possible binding of the peptides within both the proton and non-proton binding sites of ASIC3. Both sites were further investigated using molecular dynamics (MD) simulations. Analysis of the trajectories revealed potential interactions between the peptides and a set of amino acids, *i.e.* E231, E212, D252, and D358 for the proton binding site, and E247, R376, E423 and E418 for the non-proton binding site (**Fig. 6C**). **Table 1** summarizes the strength of the interactions and suggests a stronger binding of the peptides inside the non-proton binding site (strongest interaction with E418). Interestingly, the residue E247 is the only residue showing a strong difference in interaction between the two peptides, with an absolute difference of the mean interaction energy of ∼30 kcal.mol^-1^ (**Table 1**, **Fig. 6C**). This suggests a slightly different interaction on ASIC3 for the two peptides and could explain their slightly different EC_50_ values and/or their different washout kinetics. No major differences were observed between the interactions obtained from the simulations using the AlphaFold structure or the ‘post-MD’ structure (**Supplementary** Fig. 4).

**Figure 6:**
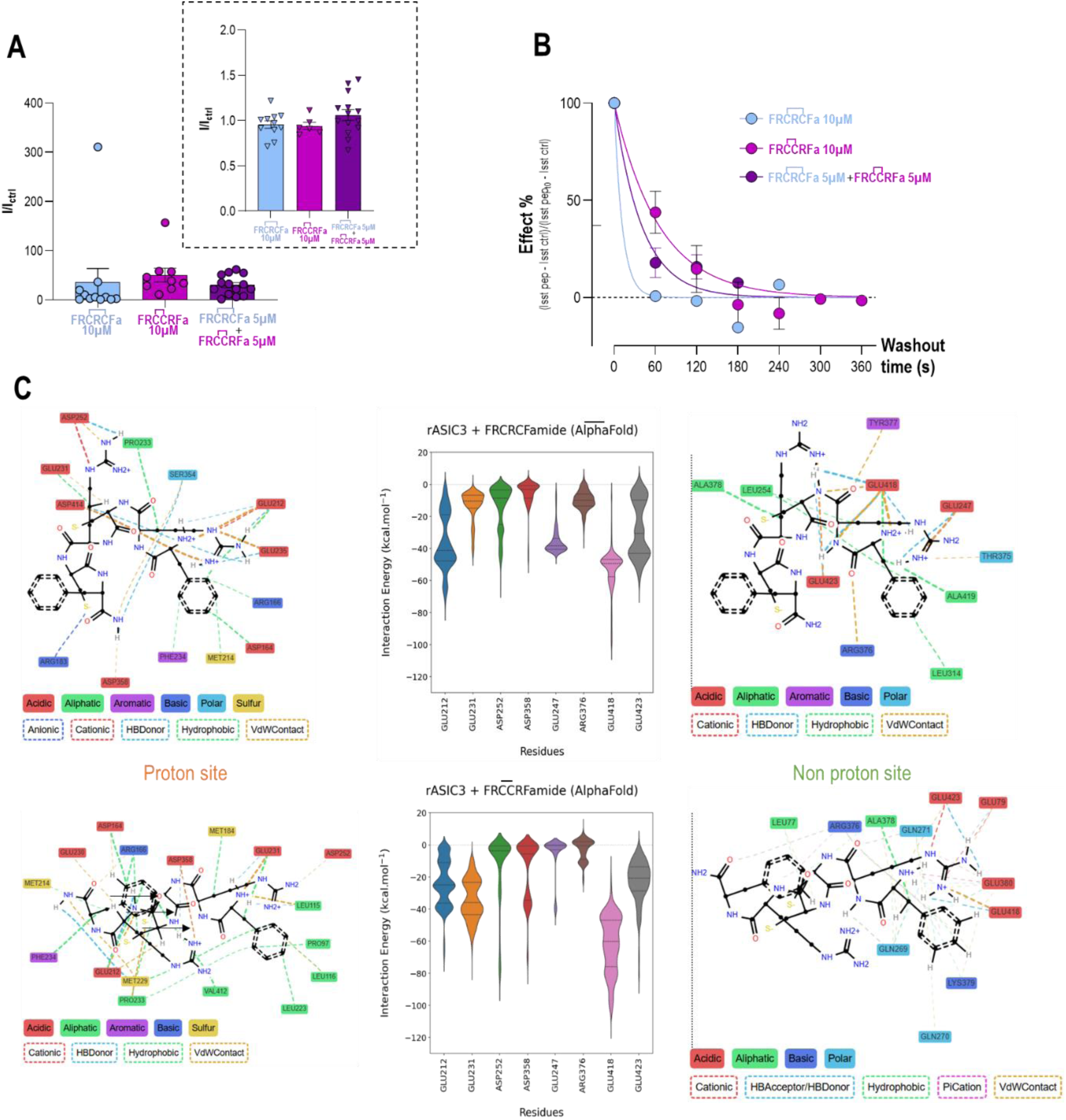
Structural elements potentially involved in the interaction of FRC̄R̄C̄Famide and FRC̄C̄RFamide with ASIC3. ***A*,** Combination of FRC̄R̄C̄Famide and FRC̄C̄RFamide at 5µM each was tested against the application of each of them alone at a concentration of 10µM, generating similar effects on pH5.5-evoked sustained current, without affecting the peak amplitude (n=9-14 cells) (***inset***). ***B*,** Washout kinetics of the effects observed in *A*, showing intermediate kinetic for the combined application of FRC̄R̄C̄Famide and FRC̄C̄RFamide at 5µM each *versus* the application of FRC̄R̄C̄Famide or FRC̄C̄RFamide alone at 10µM (same values as in Fig. 3D and 4C). ***C,*** ProLIF ligand networks for FRC̄R̄C̄Famide (upper panel) and FRC̄C̄RFamide (lower panel) interactions at the proton site (left panel) and at the non-proton site (right panel) with the violin plots of the GROMACS interaction energies for the main residues (middle panel). The thickness of the dotted lines in the ligand networks are correlated to the intensity of interaction. The dotted line in the violin plot separates the residues of the proton site to the residues of the non-proton site. The peptide backbones are shown in black.

**Table 1:** Summary of the mean interaction energy between a set of rASIC3 amino acids and both FRC̄R̄C̄Famide and FRC̄C̄RFamide peptides following the MD simulations on both the AlphaFold and the ‘post-MD’ structures. The average was taken by averaging all the interaction energies from each timestep of all 3 replicates for all 3 sites for both the proton binding sites and the non-proton binding sites using GROMACS energy ‘gmx energy’ command.

| | FRCRCFamide<br>Mean interaction energy<br>(kcal.mol<sup>-1</sup> ± std) | | FRCCRFamide<br>Mean interaction energy<br>(kcal.mol<sup>-1</sup> ± std) | |
| --- | --- | --- | --- | --- |
| rASIC3 residues | AlphaFold | Post-MD | AlphaFold | Post-MD |
| **Proton binding site** |  |  |  |  |
| E212 | -36.24 ± 17.51 | -10.16 ± 9.81 | -23.65 ± 14.42 | -25.85 ± 14.29 |
| E231 | -13.25 ± 9.51 | -34.68 ± 22.94 | -34.13 ± 12.64 | -28.01 ± 16.61 |
| D252 | -14.19 ± 13.98 | -7.68 ± 10.79 | -13.79 ± 22.02 | -0.93 ± 0.89 |
| D358 | -5.79 ± 8.68 | 10.61 ± 15.07 | -16.09 ± 19.19 | -11.67 ± 14.91 |
| **Non-proton binding site** |  |  |  |  |
| E247 | -35.73 ± 7.53 | -14.08 ± 16.30 | -5.47 ± 12.23 | -0.29 ± 0.61 |
| R376 | -9.93 ± 6.17 | -12.77 ± 10.61 | -2.93 ± 6.75 | -6.48 ± 5.35 |
| E418 | -52.76 ± 12.66 | -40.39 ± 17.23 | -61.91 ± 17.72 | -44.75 ± 14.73 |
| E423 | -27.75 ± 17.55 | -21.16 ± 21.54 | -22.79 ± 13.58 | -26.43 ± 21.76 |

Point mutations in ASIC3 of the eight amino-acids potentially involved in the predicted interactions between the peptides and the channel were next designed. All mutants were functional except for point mutation of E247. The effects of FRC̄R̄C̄Famide and FRC̄C̄RFamide peptides were tested on the different mutants (**Fig. 7**), focusing on the ability of both peptides to potentiate ASIC3 activity (**Fig. 7A-B** and see also **table 2**), but also on the washout kinetic of the FRC̄C̄RFamide effect (**Fig. 7C**). The two peptides were still able to strongly potentiate the pH5.5-evoked sustained current in the five ASIC3 mutants bearing a point mutation within the proton binding site (**Fig. 7A**), with fold-increase factors similar to those observed for the WT channel (**Table 2**). On the other hand, the sustained current potentiating effect of the two peptides was largely abolished or blunted for the E418R and E423A mutations (**Fig. 7B** and **table 2**), within the non-proton binding site (**Fig. 6B**). Both peptides had no effect on the amplitudes of the transient peak currents generated at pH5.5 by all the mutants tested (**Supplementary** Fig. 5).

**Figure 7:**
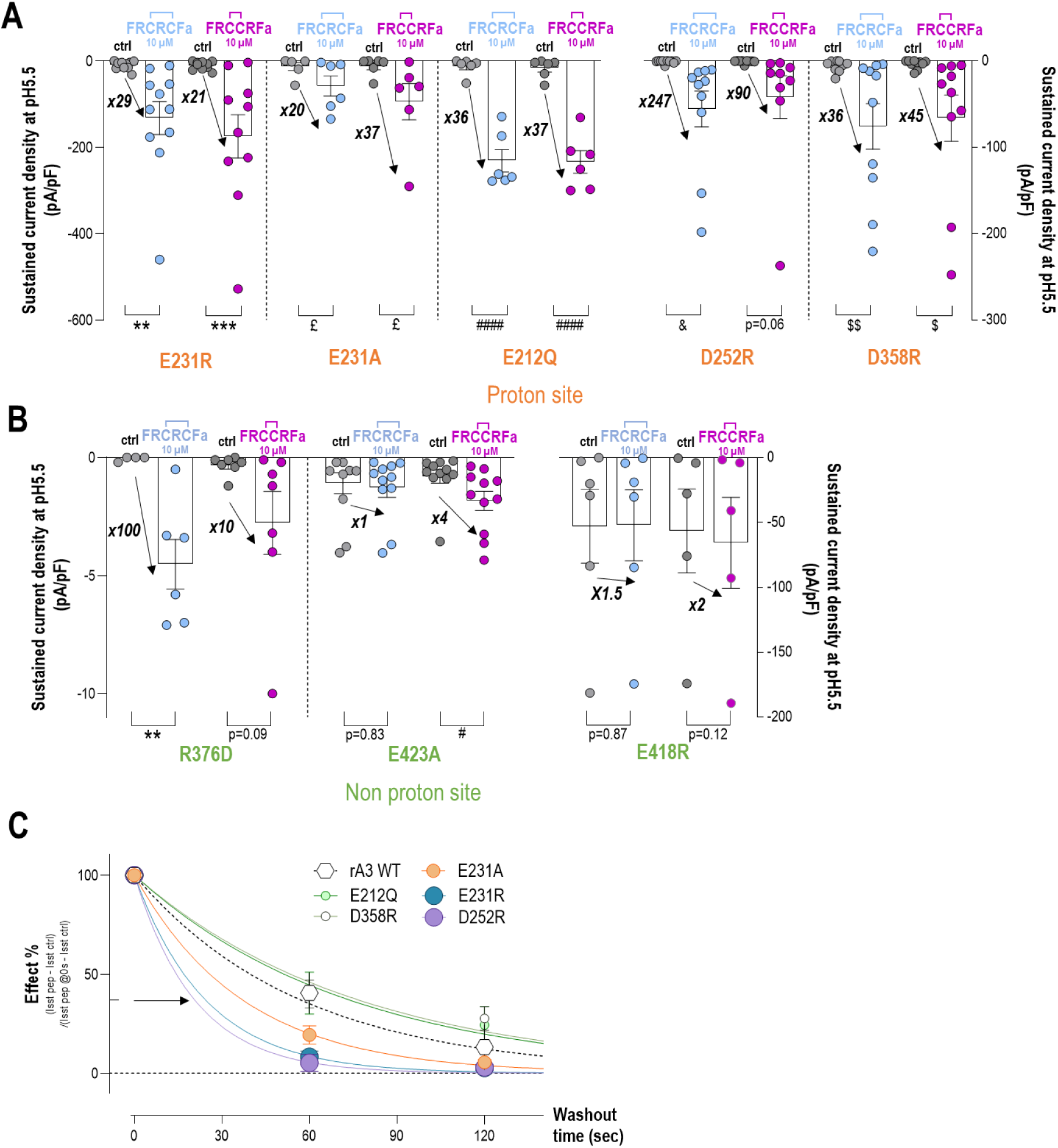
Effects of point mutations in ASIC3 of the amino-acids potentially involved in the interactions with FRC̄R̄C̄Famide and FRC̄C̄RF amide on the potentiating effect and the washout kinetic. Current densities of the pH5.5-evoked sustained currents measured in HEK293 cells transfected with different ASIC3 point mutants (holding potential -50 mV). The different bargraphs represent the current densities in pA/pF as well as the fold increase factors (arrows) induced by the two peptides. *A,* ASIC3 point mutations within the proton binding site (n=6-11, One-way ANOVA tests followed by Sidak’s multiple comparison tests). *B,* ASIC3 point mutations within the non-proton binding site (n=4-11, One-way ANOVA tests followed by Sidak’s multiple comparison tests). *C,* Slow washout kinetics of the effect of FRC̄C̄RFamide peptide on ASIC3 mutants. The washout time of the FRC̄C̄RFamide effects on six different ASIC3 mutants are represented as a function of time (WT ASIC3 channel is also represented for comparison, same data as in Fig. 3D and 4C). The washout kinetics were fitted over 120s using a one-phase exponential (see Table for the τ_off_ values of each ASIC3 mutants, n=4-18).

**Table 2:** Effect of FRC̄R̄C̄Famide and FRC̄C̄RFamide on the pH5.5-evoked sustained currents (measured 5s after current onset, see Fig. 1A) generated by wild-type (WT) and ASIC3 point mutants. The point mutations affecting the ASIC3 proton binding site are highlighted in orange, whereas those affecting the non-proton binding site are in green. The potentiating effects of the peptides is calculated as the fold increase of the sustained current (*p<0.05 and ***p<0.001, unpaired *t*-test).

|  | FRCRCFamide |  |  | FRCCRFamide |  |  |
| --- | --- | --- | --- | --- | --- | --- |
| ASIC3 type | Mean fold increase (I/I<sub>ctrl</sub>) | Significance (WT vs. mutant) | n | Mean fold increase (I/I<sub>ctrl</sub>) | Significance (WT vs. mutant) | n |
| WT | 34.08 | - | 6 | 24.22 | - | 8 |
| Proton binding site |  |  |  |  |  |  |
| E212Q | 36.1 | ns | 12 | 36.99 | ns | 9 |
| E231R | 29.37 | ns | 10 | 21.13 | ns | 9 |
| E231A | 19.86 | ns | 5 | 36.71 | ns | 6 |
| D252R | 247.36 | ns | 10 | 89.52 | ns | 9 |
| D358R | 35.54 | ns | 5 | 44.69 | ns | 5 |
| Non-proton binding site |  |  |  |  |  |  |
| R376D | 100.44 | ns | 7 | 10.2 | ns | 8 |
| E418R | 1.50 | *** (decreased) | 6 | 1.70 | ** (decreased) | 5 |
| E423A | 1.17 | *** (decreased) | 9 | 3.83 | * (decreased) | 11 |

We next studied whether the slow washout effect of FRC̄C̄RFamide on ASIC3 (**Fig. 3C**) was affected in the proton binding site mutants (**Fig. 7C**). The slow τ_off_ effect of this peptide was affected or even abolished (*i.e.*, becoming as fast as the one of FRC̄R̄C̄Famide), for the proton binding site mutants E231R, E231A and D252R (**Fig. 7C**). This may suggest that the slow washout rate of FRC̄C̄RFamide effect depends on additional peptide binding in the acid pocket, or alternatively that these mutants act indirectly by altering channel gating in different ways, which can affect the peptide washout rate from the non-proton binding site by slightly modifying conformational changes. Anyway, these data indicate a more complex mechanism of action of FRC̄C̄RFamide with effects depending directly or indirectly on the proton binding site of ASIC3, in addition to its direct binding to the non-proton binding site.

## Discussion

ASICs belong to a family of ion channels that includes the only peptide-gated channels known so far (2–4). No mammalian peptide capable of directly opening ASICs has yet been discovered, but a relatively large number of studies report that these channels have a particular sensitivity to peptides, as they are positively modulated by FMRFamide-related peptides and dynorphins (for review, see (33)). In line with these studies, a recent screening of 109 neuropeptides revealed no direct agonist of ASICs, nor any new mammalian peptide modulator of these channel, suggesting that most natural short peptides acting on ASICs are probably already known (14). The present study describes the positive modulation of ASIC3 activity by synthetic hexapeptides that were initially synthetized as NCX inhibitors 30 years ago (15). We particularly focused on two of them, a CFamide one (FRC̄R̄C̄Famide) and a RFamide one (FRC̄C̄RFamide), which show similar specificity and effects on ASIC3, but through slightly different mechanisms.

Both FRC̄R̄C̄Famide or FRC̄C̄RFamide strongly potentiate the pH-induced ASIC3 activity by increasing/slowing-down its desensitization current, generating a sustained activity, as already reported for other RFamide peptides. The two peptides show similar effect amplitude and EC_50_ (∼3µM and ∼1µM, respectively), with a good selectivity toward ASIC3, at least among ASICs, since FRC̄R̄C̄Famide had no effect on ASIC1a, ASIC1b, ASIC2a or ASIC2b homomeric channels. Such a selectivity for ASIC3 was further confirmed by experiments performed in cultured mouse DRG neurons, where the effects of FRC̄R̄C̄Famide and FRC̄C̄RFamide on native pH5.5-induced ASIC currents were lost in neurons from ASIC3^-/-^ mice. These experiments in DRG neurons also suggest that the two peptides may not only target ASIC3 homomeric, but also heteromeric channels that are largely expressed in DRG neurons (32). The small effects on the pH5.5-evoked peak current seen for native channels in DRG neurons but not for recombinant homomeric channels could also be related to the heteromeric nature of most ASIC3 channels in sensory neurons. Moreover, the two peptides do not modulate non-ASIC, TRPV1-like pH5.5-induced sustained currents (34) recorded in DRG neurons.

FRC̄R̄C̄Famide seems to act on the close state of ASIC3 channel, which is most probably also the case of FRC̄C̄RFamide, like all the other RFamide-related peptides. Indeed, all these peptides exert their effect on ASIC3 when applied extracellularly in the resting pH7.4 solution (8–10, 13). In line with this interaction mode, FRC̄R̄C̄Famide has no effect when only applied in the acid pH5.5 solution, and it has no supplemental effect when co-applied both in the resting pH7.4 plus the acid pH5.5 solutions. The potentiation of ASIC3 sustained activity was, however, strongest on currents generated by relatively more acidic pH (≤pH6.6), *i.e*., outside the ASIC3 window current (29–31), indicating a specific effect on the sustained current generated at extreme pH by particular ASIC3 structural elements (17), instead of an increase of the window current (30). If FRC̄R̄C̄Famide and FRC̄C̄RFamide share numerous similarities on the way they potentiate ASIC3 activity (almost the same EC_50_, effect amplitude and τ_on_ kinetic), we found that FRC̄C̄RFamide has a much more slower washout kinetic (τ_off_ ∼60s, five times greater than FRC̄R̄C̄Famide), indicating a poor reversibility for the effect of this peptide. RFamide peptides have already been proposed to act on ASIC3 by interacting with the non-proton binding site of the channel (12). Our data on FRC̄R̄C̄Famide and FRC̄C̄RFamide are in good agreement with such an interaction, *i.e.*, data combining FRC̄R̄C̄Famide and FRC̄C̄RFamide suggest an interaction with the same site, and effects of both peptides are lost or significantly blunted by mutations in the non-proton binding site. However, the poor reversibility of FRC̄C̄RFamide is eliminated by mutations within the proton binding site of ASIC3, which suggests a more complex mechanism of action for this peptide compared with FRC̄R̄C̄Famide. It is possible that while both peptides can interact with the non-proton binding site, the FRC̄C̄RFamide peptide could additionally interact directly or indirectly with the proton binding site, which influences its kinetic of dissociation from the non-proton binding site. Docking and molecular dynamics simulations suggest a possible direct interaction of FRC̄C̄RFamide, but also FRC̄R̄C̄Famide, with the proton binding site, although weaker than for the non-proton binding site. However, only FRC̄C̄RFamide has a slow washout kinetic, and FRC̄R̄C̄Famide has to be linearized to display a slow one. In addition, although those two peptides are supposed to bind into the acidic pocket (computational modelling), none of them affects the transient peak current, which would be expected from the molecular changes associated with proton activation of ASICs such as the collapse of the acidic pocket (35). Although binding of neuropeptides into the acidic pocket of ASICs has already been described for dynorphin and ASIC1a (36), it is more likely that all the RFamide and CFamide peptides described here only bind to the non-proton binding site of ASIC3. Conformational constraints for some of them, as suggested for instance by modelling data showing a strong difference in interaction between FRC̄C̄RFamide and FRC̄R̄C̄Famide peptides for residue E247, lead to a slow washout kinetic, which can be overcome either by cyclization (as for FRC̄R̄C̄Famide, but not FRC̄C̄RFamide) or by indirect gating-related effects associated with the proton binding site when this site is mutated.

In summary, the present study identifies a set of short amidated hexapeptides as new ASIC3 ligands, with strong potentiating effects of the channel activity. Focusing our investigations on FRC̄R̄C̄Famide and FRC̄C̄RFamide led us to identify similarities in these peptide effects, in line with the effects already reported for other peptides of the RFamide family, including a possible interaction with the non-proton binding site. An important point of our study could be the identification of some short amidated peptides with long lasting effects (poor reversibility of FRC̄C̄RFamide, FRCCRFamide and FRCRCFamide) providing new potential pharmacological tools, and suggesting a more complex mechanism of action than the one previously describes for other RFamide peptides, which could also involve a regulation through the acidic pocket of ASIC3. This raises the possibility that RFamide peptides may also bind or secondarily interact with the acidic pocket of ASIC channels, which is in line with recent data showing that the ligand-binding pocket in the HyNaC channels from the cnidarian *Hydra magnipapillata* that are activated by Hydra RFamide peptides is localized in the same region as the acidic pocket of ASICs (37).

## Acknowledgement

We thank Drs A. Baron, S. Diochot and J. Noel for helpful discussions, V. Friend and V. Thieffin for technical support, and J. Thoral for secretarial assistance. This work was supported by the Centre National de la Recherche Scientifique (CNRS), the Institut National de la Santé et de la Recherche Médicale (Inserm), the Association Française contre les Myopathies (AFM grant #23731), the Agence Nationale de la Recherche (ANR-17-CE16-0018 and ANR-22-CE16-0006), the Institut ANALGESIA/SFETD (Société Française d’Etude et de Traitement de la Douleur), the LabEx ICST (ANR-11-LABX-0015-01), the Fondation pour la Recherche Médicale (FRM grant #FDT202404018117), the University Côte d’Azur (UniCA) and the French government, through the UCA^JEDI^ Investments in the Future project managed by the ANR with the reference number ANR-15-IDEX-01.

## Author contributions

M.T., M.M., H.L.D.S. designed, performed and analyzed electrophysiological recordings. C.V. and P.Z. did the computational modelling part of the work. M.S. designed and performed mutagenesis experiments. E.L. and E.B. helped in scientific design and critical reading of the manuscript. E.D. conceived the project, did patch clamp experiments, data analysis and wrote the paper with the input of all authors.

---

**Supplementary Figure 1:**
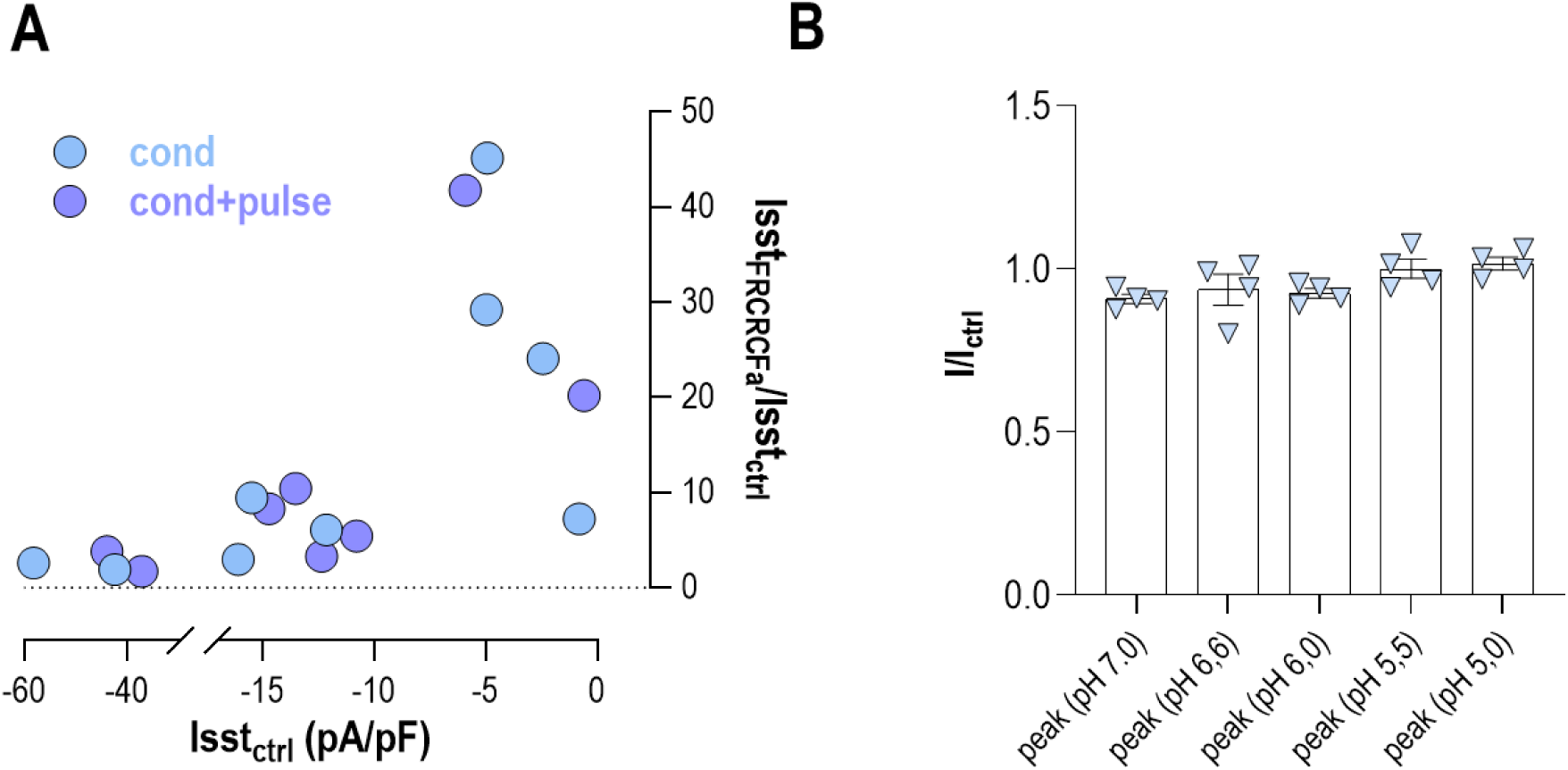
Application of FRC̄R̄C̄Famide (1µM) does not change peak amplitude, regardless of pH stimulation. (**A**) Negative correlation between the ASIC3 sustained current fold-increase factor induced by FRC̄R̄C̄Fa and the basal amplitude of this sustained current (Isst_ctrl_, Spearman correlation with r=-0.74, p=0.0010, data from conditioning (cond) and conditioning and pulse (cond+pulse) conditions, (n=17). (**B**) FRC̄R̄C̄Famide at 1µM was applied 30 seconds before 10 seconds stimulations at pH 7.0, 6.6, 6.0, 5.5, 5.0, and the peak of pH-evoked ASIC3 was unchanged (n=4).

**Supplementary Figure 2:**
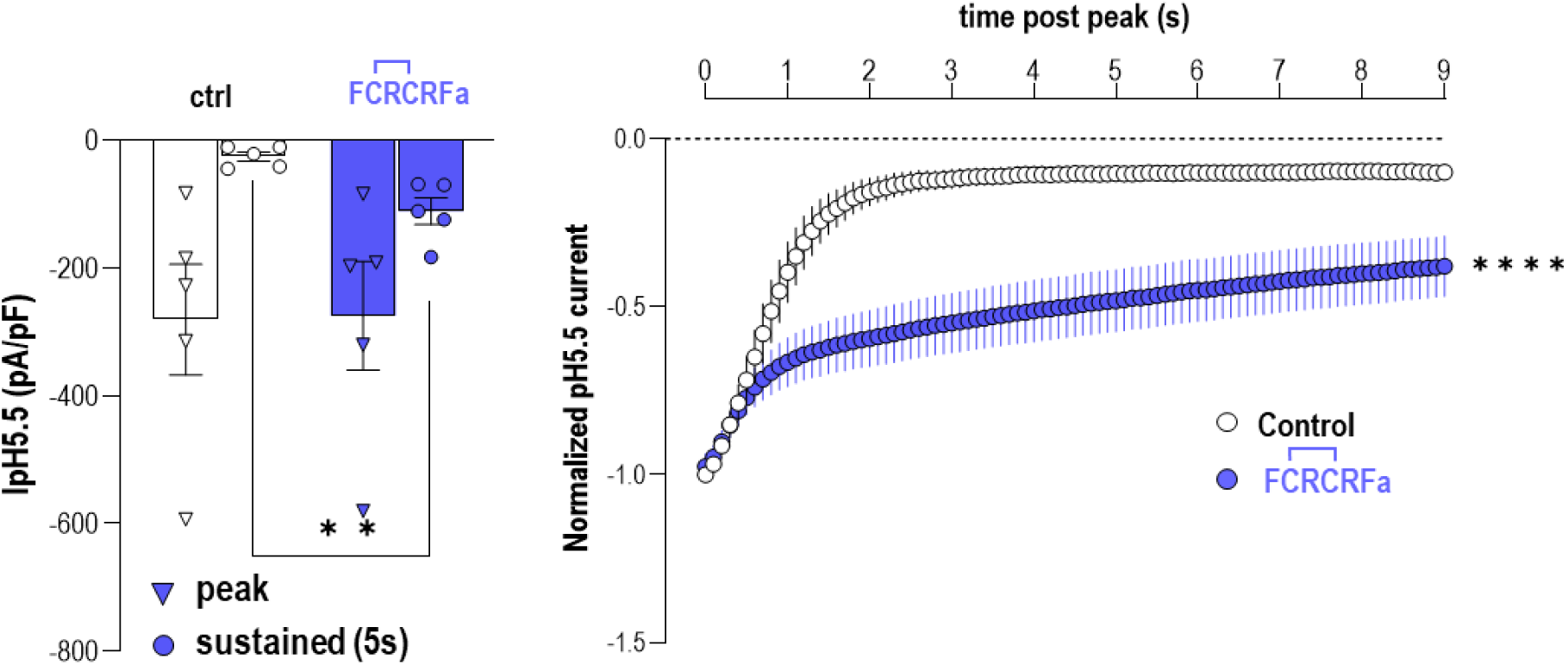
FC̄R̄C̄RFamide peptide potentiate ASIC3 sustained current. Analysis of the effect of FC̄R̄C̄RFamide peptide (10µM in the pH7.4 resting solution) on rat ASIC3 peak and sustained current densities elicited at pH5.5 before and after extracellular applications of peptide (n=5 ** p<0.01, mixed-effect analysis followed by Dunnett’s multiple comparisons test). Current amplitudes were measured every 100ms during the 9s following the peak, and normalized to the control peak amplitude before application of each peptide (n=5, two-way ANOVA with ****p<0.0001).

**Supplementary Figure 3:**
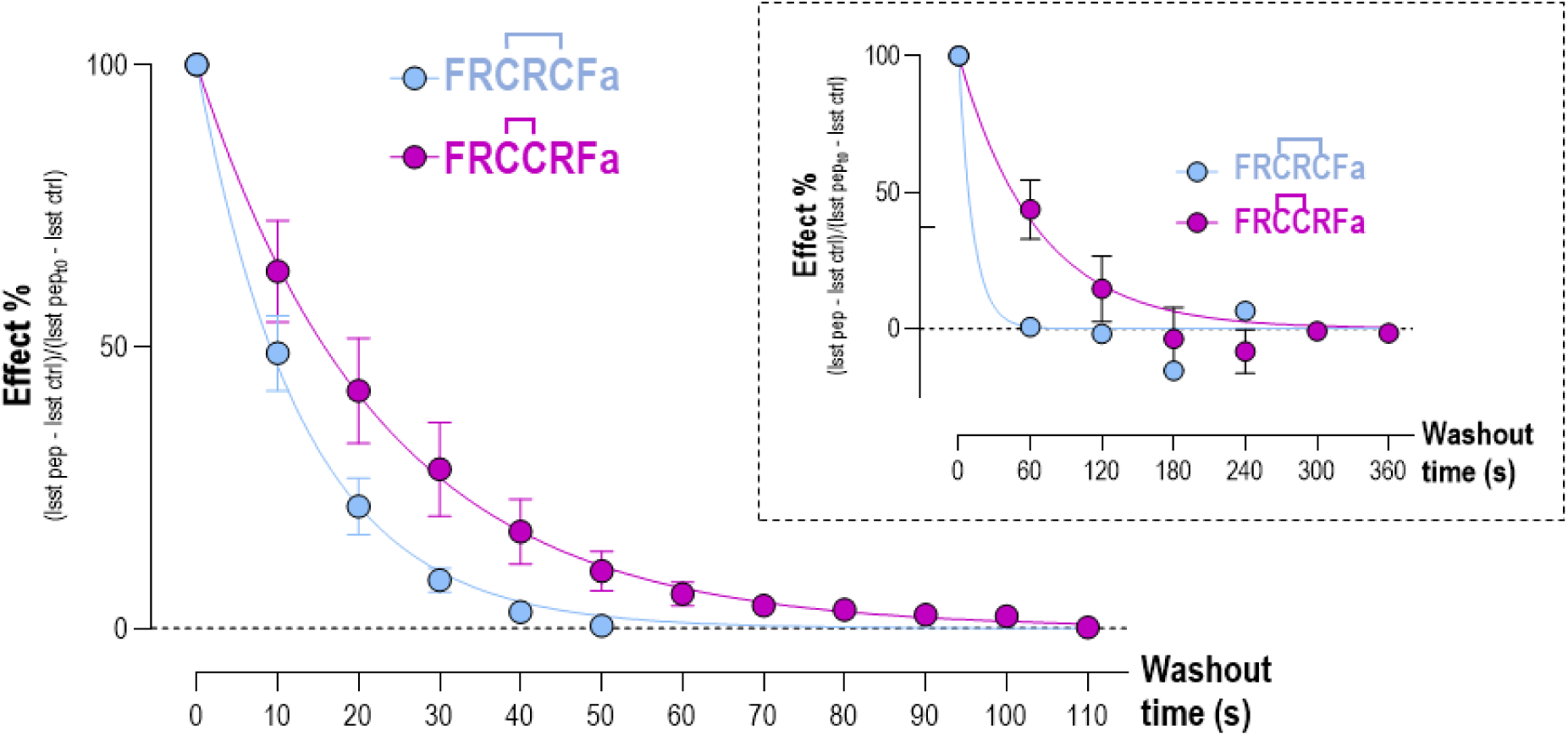
Increasing the frequency of pH5.5 stimulations (every 10s) following FRC̄C̄RFamide application accelerate its washout kinetic, but not that of FRC̄R̄C̄F amide. The washout kinetics were fitted over 60s using a one-phase decay exponential (τ_off_= 13.1s for FRC̄R̄C̄Famide and τ_off_= 22.7s for FRC̄C̄RFamide, n=6-7). For comparison, the *inset* illustrates the washout kinetics of FRC̄R̄C̄Famide and FRC̄C̄RFamide already shown in Fig. 3D and obtained using a low-frequency washout protocol (pH5.5 stimulations every 1 min, (τ_off_=11.9s and 65.5s, respectively). The τ_off_ for FRC̄R̄C̄Famide remains unchanged in the two washout protocols, but not that of FRC̄C̄RFamide.

**Supplementary Figure 4:**
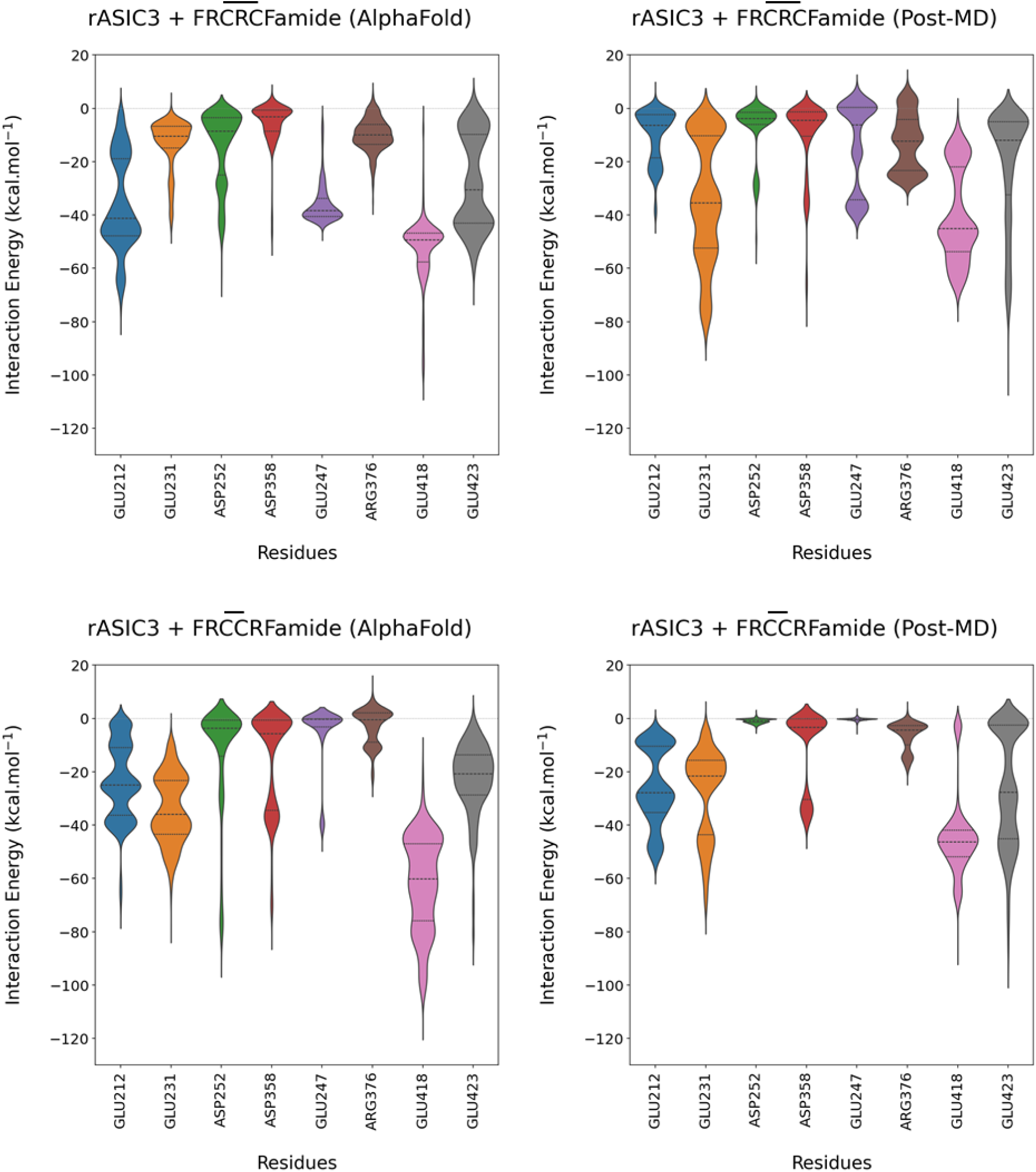
Comparison of interaction energies between simulations using the AlphaFold3 structure (left panels) or the post-MD structure (right panels) as input. Top two panels show the differences between the interaction energies of rat ASIC3 residues in the proton (E212, E231, D252, and D3358) and non-proton (E247, R376, E418, and E423) binding sites with the FRC̄R̄C̄Famide peptide. The average difference of the means is 12.79 kcal.mol^-1^ (with the main difference of 26.08 kcal.mol^-1^ observed for E212). Bottom two panels show the differences between the interaction energies of rASIC3 residues of the proton and non-proton sites with the FRC̄C̄RFamide peptide. The average difference of the means is 6.89 kcal.mol^-1^ (with the main difference of 17.16 kcal.mol^-1^ observed for E418).

**Supplementary Figure 5:**
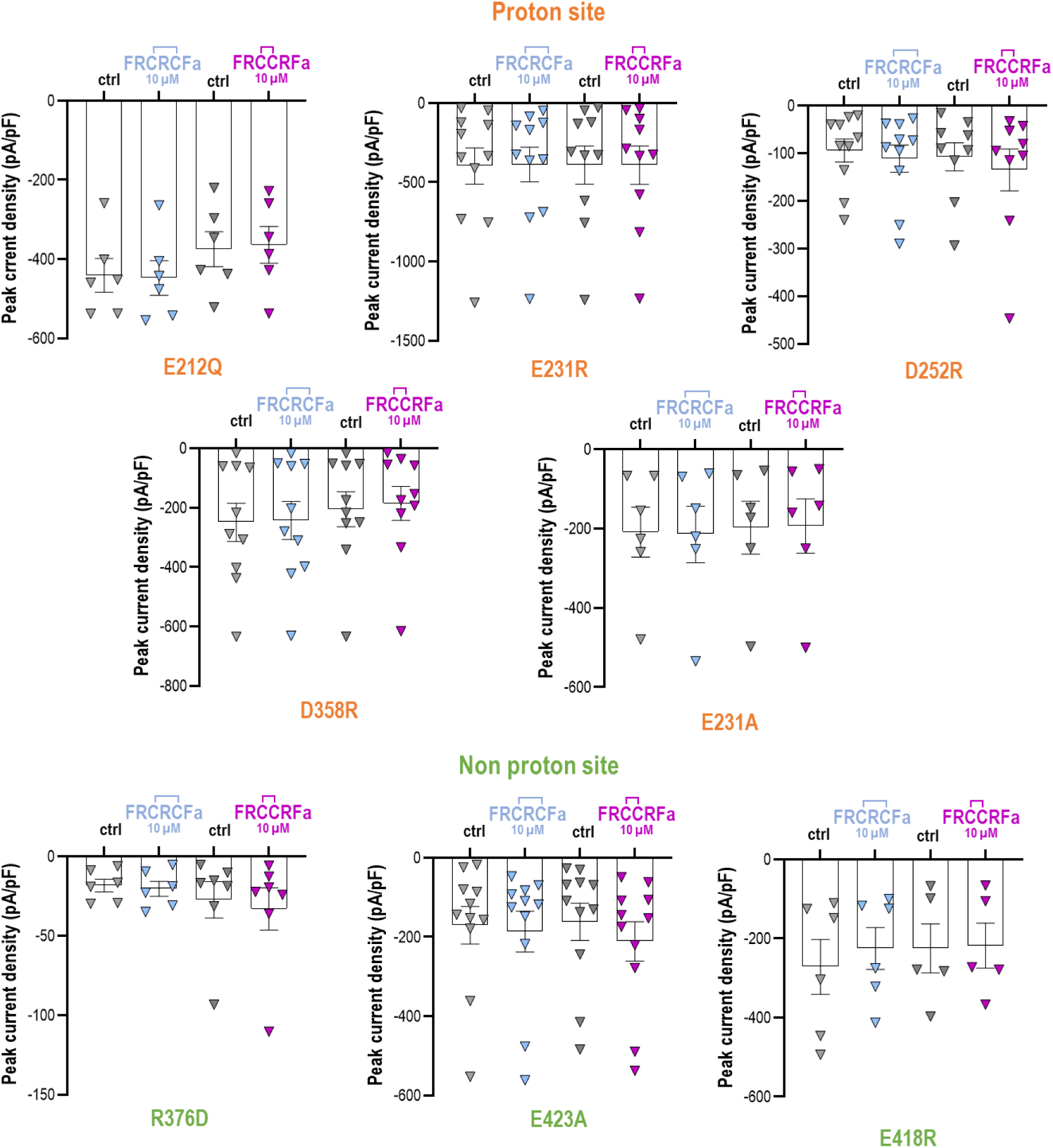
Effects of point mutations in ASIC3 of the amino-acids potentially involved in the interactions with FRC̄R̄C̄Famide and FRC̄C̄RFamide on the peak current densities (pA/pF). Current were recorded at -50mV and were elicited by pH5.5 drops. Mutations affecting the proton and non-proton binding sites of ASIC are indicating in orange and green, respectively. Peptides were applied in the pH7.4 resting solution for 30s before the pH5.5 drops (no significant differences were observed, One-way ANOVA followed by Sidak’s multiple comparison tests).

